# The immunobiology of persistent intestinal infection by *Campylobacter jejuni* in the chicken

**DOI:** 10.1101/2024.08.23.609314

**Authors:** Rachel Gilroy, Amy Wedley, Sue Jopson, Judith Hall, Paul Wigley

**Affiliations:** Institute of Infection, Veterinary and Ecological Sciences, University of Liverpool, Leahurst Campus, Neston, CH64 7TE UK; Biosciences Institute, Newcastle University, Newcastle upon Tyne, NE2 4HH; Bristol Veterinary School University of Bristol, Langford, Bristol, BS40 5DU UK

## Abstract

Infection of the intestinal tract of the chicken with *Campylobacter jejuni* is frequently prolonged with limited immune clearance from the caeca, the main site of colonisation. Previously it has been shown early infection of broiler chickens leads to a pro-inflammatory response, followed by regulatory and Th17 responses along with adaptive Th1 and Th2 cytokine responses. Here we show that infection up to 28 days post challenge leads to prolonged expression of IL-4 and high titre IgM and IgY serum antibody responses to *C. jeuni*, along with increased levels of total IgA in the gut of infected birds. Whilst early pro-inflammatory cytokine and chemokine responses largely wane, expression of both IL-10 and IL-17A remain high in the colonised caeca or caecal tonsils. Based on this and previous studies we hypothesize that the nature of the immune response to *C.jeuni* infection is one that allows persistence in the gut, but limits inflammation and invasive infection from the gut through maintaining the intestinal barrier and producing secretory antibody. This protects the birds from campylobacteriosis, but the persistence in the gut for the production lifetime of chickens remains an unsolved public health issue.

## Introduction

*Campylobacter* is the most frequent bacterial cause of foodborne gastrointestinal disease with the majority of human campylobacteriosis caused by *Campylobacter jejuni*. Although multiple sources of *C. jejuni* infection ranging from red meat to the natural environment have been described, most cases of campylobacteriosis are associated with the consumption of contaminated poultry meat. The ability of *C*.

*jejuni* to colonise the intestinal tract of chickens, to spread rapidly and flocks and persist in sufficient numbers through slaughter and processing result in the frequent contamination of poultry meat in retail and catering(1). Although a range of control measures to reduce *C. jejuni* in chicken production have been proposed and implemented with limited success(2), current opinion is that control through vaccination or feed-based interventions are most likely to be successful. For either of these potential interventions an understanding of both host and pathogen processes that underpin intestinal colonisation and persistence, including a fuller understanding of the longer-term immune response are needed.

*C. jejuni* has until recently been considered ‘merely a commensal’ of the chicken gut. It is now clear that infection causes initial inflammation and changes to gut permeability leading to low level pathology associated with poor gut health and reduced welfare in broiler chickens(3). This inflammatory damage is associated with expression of key inflammatory cytokines and CXC chemokines along with lower expression of regulatory cytokines such as IL-10(4). As well as activation of innate responses, *C. jejuni* elicits a strong adaptive response with high levels of specific antibody along with some evidence of cellular responses(4, 5). Whilst both vaccination and infection lead to antibody responses there is limited and somewhat conflicting evidence of their role in protection or clearance from the main site of colonisation, the ceca(6). Depletion of antibody responses by neonatal treatment with cyclophosphamide (chemical bursectomy) has limited effect in persistence of infection, though antibody does appear to reduce *C. jejuni* levels in the ileum(7). The general absence of clearance has led to the suggestion the response to *Campylobacter* is ‘tolerogenic’ where invasive infection is prevented but persistence in the gut allowed(8). Modelling of cytokine networks following infection supports this idea as initial inflammation is largely controlled and that Th17 responses are a key response(9). Although IL17/Th17 responses are often associated with inflammatory conditions, they are also key sentinels in maintaining gut integrity through maintenance of epithelial tight junctions and increasing expression of antimicrobial peptides at the gut epithelium(10, 11). We suggested previously that such responses may drive the longer-term balance between immunity and colonisation(9). Here we show mucosal expression of key cytokines in gut mucosal tissue and specific antibody production which suggest an immunological basis by which *C. jejuni* persists in the chicken intestinal tract.

## Materials and Methods

### Experimental animals

Experimental infection trials were carried out at the University of Liverpool poultry unit (Liverpool, UK) in accordance with United Kingdom legislation governing experimental animals, Animals (Scientific Procedures) Act 1986. All work was conducted under project licences PPL 40/3652 (Experiment 1) and P999B8C93 (Experiment 2) was approved by the University of Liverpool ethical review process prior to the award of the licence by the UK government. All animals held at the site were checked a minimum of twice daily to ensure individual and flock animal health and welfare. All *in vivo* experiments used day-old broiler chicks (Ross 308) of mixed sex obtained from a local commercial hatchery. Ross 308 remains the most commonly reared broiler breed within the UK, justifying its use within these experimental models. All chicks were transported directly from the hatchery environment to the experimental unit and observed for any potential adverse indications.

Chicks were maintained according to treatment group in separate experimental rooms within floor pens at a stocking density in accordance with Home Office Code of practice recommendations. All rooms were supplied with filtered air supply with regular air changes, while groups intended for experimental infection protocols were housed in rooms with lobbied entry and additional dedicated protective clothing and boots. All animals were housed in conditions previously described (4).

Birds were given *ad libitum* access to water and a pelleted vegetable protein-based diet (SDS, Witham, Essex, UK). Feeders and drinkers were provided at a level of 1 per 15 birds. Room temperature was kept at a temperature of 30°C before being reduced to 20°C when the birds were three weeks of age. To limit welfare problems associated with wet litter and to limit within-group retransmission, litter was changed, and pens were cleaned once every four days.

At 14 d.p.h (days post hatch), *Campylobacter* negative status was confirmed for all birds prior to experimental infection through cloacal swabbing. Swabs were subsequently streaked onto *Campylobacter*-selective blood-free agar (modified charcoal-cefoperazone-deoxycholate agar [mCCDA]) supplemented with *Campylobacter* enrichment supplement (SV59; Mast Group Ltd, Bootle, Merseyside, UK) before incubating at 41.5 ^°^C for 48-hours in microaerobic conditions.

### *Campylobacter jejuni* challenge strain

*C. jejuni* M1 sequence type (ST)137 (clonal complex [CC] 45) well characterized for its genetics, virulence and ability to colonise chickens was used in this study (12) *C. jejuni* Mi stock was maintained at - 80 °C on Microbank™ beads (ProLab Diagnostics, Cheshire, UK) until use. Briefly, bacteria were grown on Columbia blood agar (CAB) (Lab M Ltd., Heywood, Lancashire, United Kingdom) supplemented with 5% defibrinated horse blood (Oxoid, Basingstoke, Hampshire, United Kingdom) at 41.5 °C for 48-hours under microaerobic conditions (80 % N_2_, 12 % CO_2_, 5 % O_2_ and 3 % H_2_).

### Preparation of inoculum

*C. jejuni* M1 challenge inoculum was grown in Mueller Hinton broth (MHB) (Lab M Ltd., Heywood, Lancashire, United Kingdom) as previously described (13)

### Experimental design – Experiment 1

Age-matched, 1 d.p.h mixed sex Ross 308 chicks (*n* = 87) were introduced to the University of Liverpool high-biosecurity poultry unit under housing conditions described previously. On point of entry, the chicks were randomly assigned to one of two groups: Group 1 (*n*=57) or Group 2 (*n* = 30).

Prior to infection, at 14 d.p.h, all animals were confirmed to have *Campylobacter* negative status as previously described. At 21 d.p.h, all birds within the *C. jejuni* Group 1 (*n* = 57) were orally infected with 0.2 mL 10^6^ CFU/mL *C. jejuni* in MHB via oral gavage. All birds in Group 2 (*n* = 30) were given 0.2 mL sterile MHB via oral gavage. Preparation of inoculum and infection protocols were conducted as described previously. From this point, experimental groups 1 and 2 were named *C. jejuni* infected and non-infected control groups respectively.

At 23 (2 days post infection [d.p.i]), 28 (7 d.p.i), 35 (14 d.p.i), 42 (21 d.p.i) and 49 (28 d.p.i) days post hatch, randomly selected birds were culled via cervical dislocation from the *C. jejuni* infected group (see **Error! Reference source not found**.) and non-infected trial group (n = 6). Blood samples were collected via cardiac puncture immediately post-cull, before samples of splenic & liver tissues and caecal & ileal content were aseptically collected. An additional ileal tissue section was collected, and ileal content removed for subsequent gut wash processing.

### Experimental design – Experiment 2

Age-matched, 1 d.p.h mixed sex Ross 308 chicks (n = 53) were introduced to the University of Liverpool high-biosecurity poultry unit under housing conditions described previously. On point of entry, the chicks were randomly assigned to one of two groups: Group 1 (n=26) or Group 2 (n = 27).

Prior to infection, at 14 d.p.h, all animals were confirmed to have *Campylobacter* negative status as previously described. At 21 d.p.h, all birds within the *C. jejuni* Group 1 (n = 26) were orally infected with 0.2 mL 106 CFU/mL *C. jejuni* in MHB via oral gavage. All birds in Group 2 (n = 27) were given 0.2 mL sterile MHB via oral gavage.

At 23 (2 days post infection [d.p.i]), 28 (7 d.p.i), 35 (14 d.p.i) and 42 (21 d.p.i) days post hatch, a pre-defined number of randomly selected birds from each group were culled via cervical dislocation (**Error! Reference source not found**.). Blood samples were collected via cardiac puncture immediately post-cull, before samples of splenic & liver tissues and caecal & ileal content were aseptically collected.

### Quantitative bacteriology

Tissue and gut content samples were enumerated on mCCDA agar as described previously (13)

### Determining serum IgY and IgM specific antibody responses

Chicken serum IgM and IgY responses to *C. jejuni* were determined by ELISA using blood samples collected at post-mortem according to protocols previously described (7).

### Measurement of secretory IgA

As a means of determining secretory IgA levels within the ileum, a 25 cm section of ileal tissue (without ileal content) was aseptically collected at *post mortem* examintion. Ileal sections were flushed with 10 mL of sterile 1 x PBS while the tissue was manually massaged. The flush was collected and centrifuged for 10 minutes at 500 x *g* before aseptic collection of the supernatant with a pipette. All processed samples were stored at – 20 °C until further processing.

Quantification of secretory IgA within the processed samples was then conducted using an IgA Chicken ELISA Kit according to manufacturer’s instructions (ab15, Abcam®, Cambridge, UK). Briefly, 100 µl of pre-prepared standard solutions (concentrations ranging from 12.5 – 400 ng/mL) including blank control (consisting only of provided diluent solution) were added in duplicate into pre-designated wells of a provided 96-well plate. Gut wash samples were diluted 1:5000 in 1 x diluent solution provided and added to wells of the provided 96-well plate in duplicate.

Plates were incubated for 20-minutes at RT before being washed 4 times with 1 x wash buffer (provided). 100 µl of 1 x enzyme-antibody conjugate (provided) was added to each well and incubated for 20 minutes at RT in the dark. Plates were washed 4 times with 1 x wash buffer (provided) and 100 µl of TMB substrate solution (provided) was added to each well before incubation in the dark at room temperature for 10-minutes. Absorbance was determined using a microplate reader at 405 nm and a standard curve plotted to interpolate total secretory IgA concentrations in ng/mL.

### RNA extraction

Tissue samples of spleen, caeca, caecal tonsil and ileum were collected from all infected and control birds of experiment 1 and stored in 1 mL RNAlater at -20°C until further processing (Sigma, Poole, Dorset, UK). Total RNA was extracted from 20 – 30 mg of all tissue samples using a RNeasy minikit (Qiagen, West sussex, United Kingdom) according to manufactures instructions. Prior to extraction protocols, Buffer RLT solution (provided) was first supplemented with 10 µl of β-mercaptoethanol (Sigma, Poole, Dorset, UK) per 1 mL Buffer RLT. All tissues were disrupted using a TissueLyser (Qiagen, West Sussex, UK) at a frequency of 10,000/S for 10 minutes, with the addition of one stainless steel metal bead per sample. Isolated RNA was eluted into 50 µl of RNase-free water and stored at -80 °C until processing. Total RNA yield per sample was determined using a Nanodrop (ND-1000) spectrophotometer, with samples showing low yield being re-extracted.

### Cytokine gene expression

mRNA expressional changes for the cytokines interleukin-1β (IL-1β), IL-4, IL-6, IL-10, IL-17A, TGFβ_4_ and the chemokine ligand CXCLi2 were measured in these tissue samples by real-time quantitative reverse-transcription PCR (qRT-PCR) using a Rotor-Gene Q version 2.3.1.49 (Qiagen, West Sussex, UK) as previously described by (Humphrey et al 2014).

### Expression of Mucin 2

mRNA expressional changes in the AMP Mucin2 were measured in caecal tissue biopsy samples by qRT-PCR using a Rotor-gene Q version 2.3.1.49 (Qiagen, UK) as described previously (Humphrey et al, 2014). Primer sequences used are listed in **Error! Reference source not found**. alongside the respective threshold value. For each reaction, β-actin rRNA (ACTB) was used as the housekeeping gene to normalise mRNA levels between samples. Threshold values for each gene transcript were determined at 10 % of the curve plateau of known *Campylobacter*-positive control samples. All RNA samples were first diluted 1:10 using RNase-free water to obtain desired concentration per sample.

One-step RT-PCR was performed using the Precision®PLUS OneStep RT-qPCR Master Mix premixed with SYBRgreen (PrimerDesign, Camberley, UK) to a final reaction volume of 20 µl according to manufacturers’ instructions. All reactions contained 2 µl of total RNA (at a total concentration of 25 ng), 10 µl of the Precision®PLUS OneStep RT-qPCR Master Mix premixed with SYBRgreen, 0.6 µl forward primer (at 6pmols), 0.6 µl reverse primer (at 6pmols), and 6.8 µl of RNase-free water.

Each sample was assessed in triplicate, with no-template control samples containing RNase-free water in place of total RNA being used per run. All reactions were conducted according to the following reaction profile listed in Table 1

**Table 1.** Details of qRT-PCR amplification conditions using SYBR Green PCR.

|  | Step | Time | Temperature |
| --- | --- | --- | --- |
|  | Reverse<br>Transcription | 10 minutes | 55°C |
|  | Enzyme<br>Activation | 2 minutes | 95°C |
| ***Cycling x 40*** | Denaturation | 10 seconds | 95°C |
|  | Data Collection | 60 seconds | 60°C |
|  | Melt Curve |  | 50°C - 99°C |

### Analysis of gene expression

For each expression triplicate per sample, an average Ct value was calculated based on threshold value used per gene of interest. All expression values for target genes were determined using 2-^ΔΔ^C_t_ methodologies. Firstly, gene of interest (GOI) C_t_ was determined relative to that of the housekeeping 28S rRNA (^Δ^C_t_). ^Δ^C_t_ values given for samples from *C. jejuni* infected birds were then normalised against those of uninfected control animals to give final readings as relative fold changes (2-^ΔΔ^C_t_).

To determine statistical significance of variations in transcript expression between control and infected samples, pairwise comparisons of 40 - ^Δ^C_t_ was performed using Mann Whitney-U analysis, with statistical significance set at p < 0.05.

### Statistical analysis

Statistical analysis was performed using GraphPad Prism version 7.00 for Mac OS X (GraphPad Software Inc., San Diego, USA). Prior to further statistical analysis, all data was first assessed for distribution normality using D’Agostino-Pearson omnibus normality testing. Pairwise treatment group comparisons of normally distributed data sets (p > 0.05) were conducted using an Unpaired *t*-test and described using data mean and standard deviation values (SD). Pairwise treatment group comparisons of non-normally distributed data sets (p < 0.05) were conducted using a Mann Whitney-U test and described using data median and Interquartile range (IQR). Statistical significance was determined using a p < 0.05 threshold. For statistical comparisons assessing more than two distinct groups, Kruskal-Wallis testing was used with a p <0.05 threshold.

## Results

### Colonisation of the chicken gut with *C. jejuni*

Infection with *C. jejuni* M1 leads to persistent colonisation of the caeca in broiler chickens. As expected in each experiment *C. jejuni* MI colonised the caeca at a high level and for the duration of the experiment (Figure 1)

**Figure 1.**
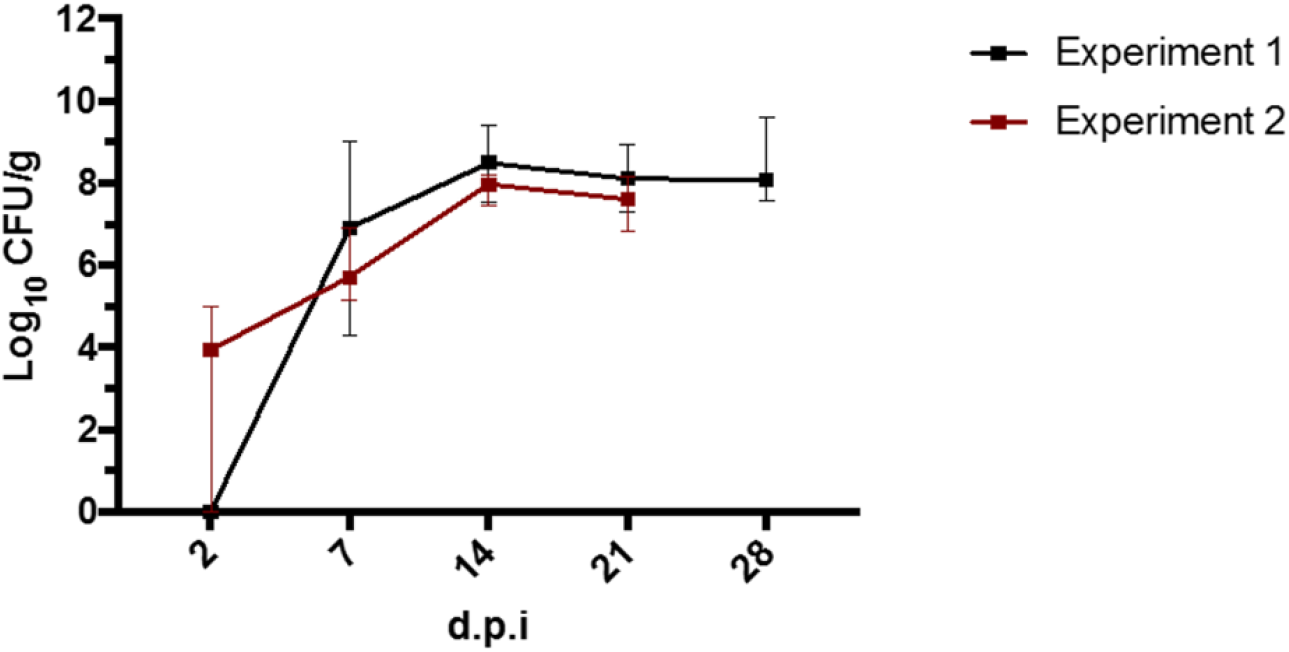
Oral challenge with *C. jejuni* M1 leads to persistent colonisation of the caeca. Mean caecal load of *C. jejuni* M1(± SEM) in Ross 308 broiler chickens at days post infection (dpi) following challenge at 21 days-of-age

### IIL-10 and IL-17A are expressed in the caeca during persistent caecal colonisation

Initial expression of cytokines at 2 days (not shown) and 7 days post challenge (Figure 2) followed those previously described with expression of pro-inflammatory cytokines(4, 9). From 14 days post infection cytokine there is significant expression of IL-17A (P<0.05) and TGF-β4 (P<0.01) in the caeca. Expression of TGF-β4 remains significantly upregulated at 28 days post challenge along with IL-10 and highly significant expression of IL-17A. In contrast, expression of pro-inflammatory and Th1 and Th2 associated cytokines are largely both caeca and caecal tonsil at 21 days (data not shown) and 28 days post infection (Figure 4). In the caecal tonsil there is consistent upregulation of both IL-4 and IL-10 at all time points.

**Figure 2.**
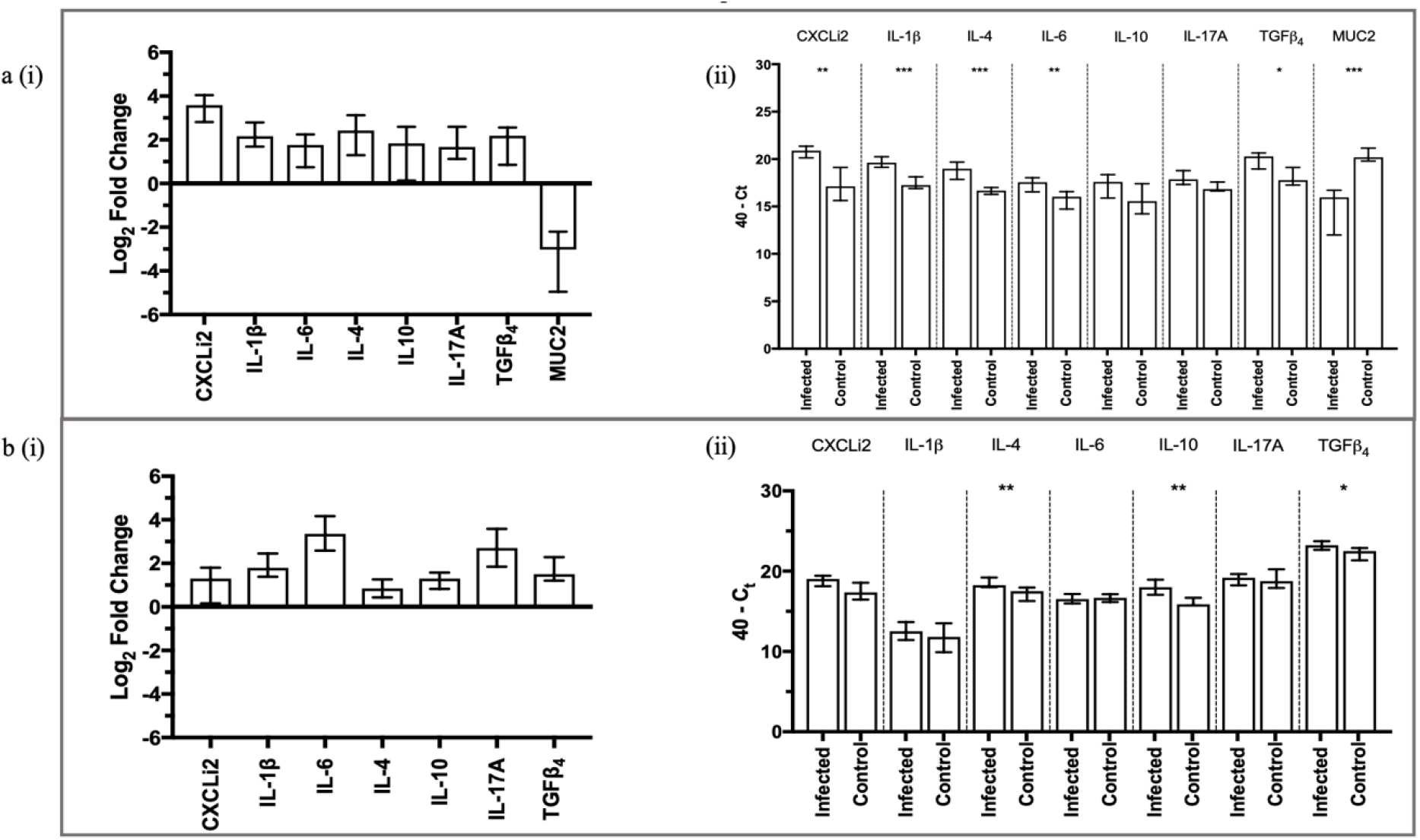
Cytokine and Mucin expression seven days post infection. Relative expression of infected v control (i) and 40 - ^Δ^Ct (ii) of cytokine, chemokine and AMP transcripts within caecal tissue (a) and caecal tonsil tissue (b) of experimental chickens according to *C. jejuni* challenge status at 7 days post infection. Error bars represent IQR of the median value. Statistical significance was determined on 40 – ^Δ^Ct data using Mann Whitney-U analysis with levels of significance given as * p < 0.05, **p < 0.01, ***p < 0.001

**Figure 3.**
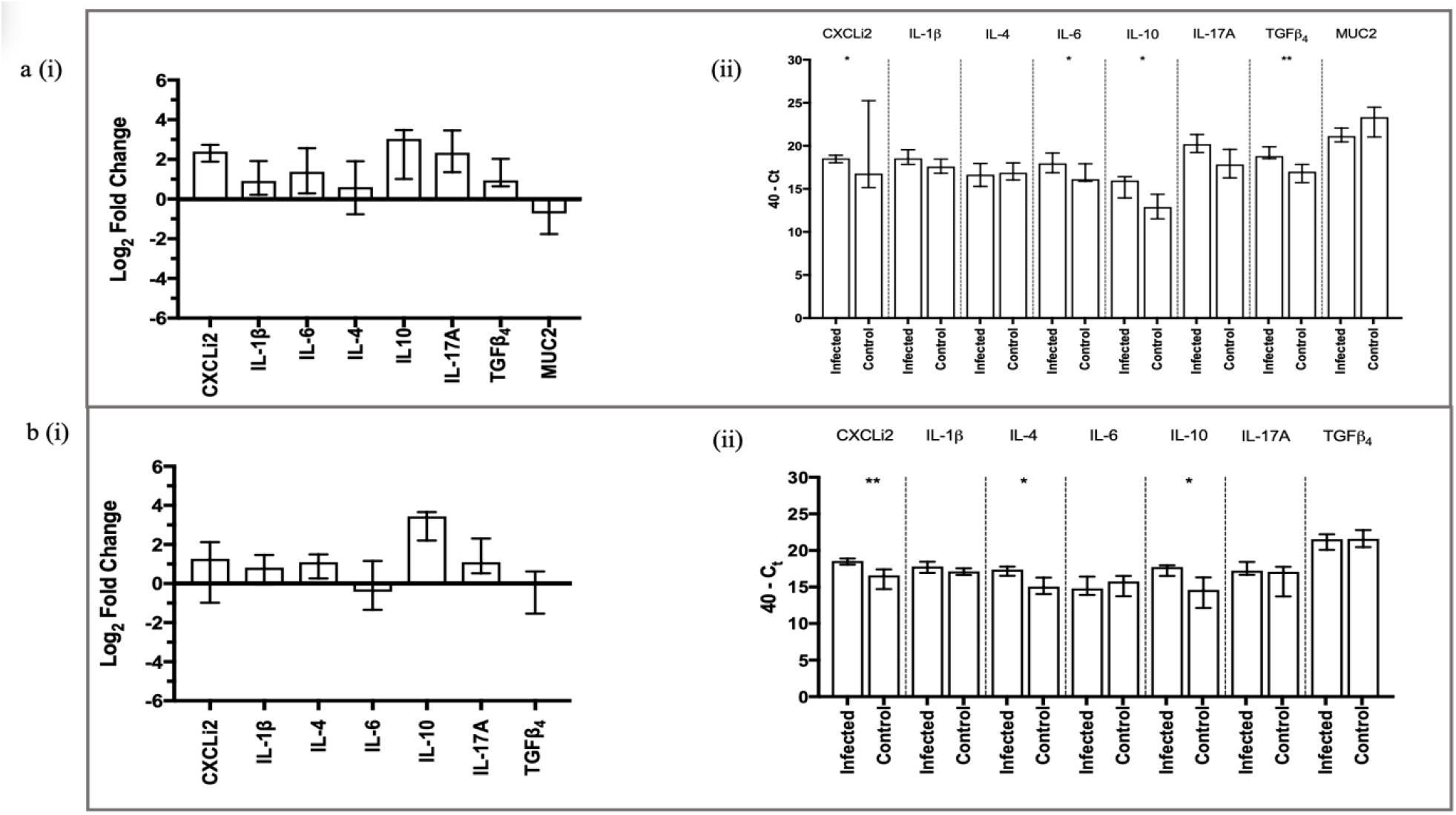
Cytokine and Mucin expression 14 days post infection. Relative expression of infected v control (i) and 40 - ^Δ^Ct (ii) of cytokine, chemokine and AMP transcripts within caecal tissue (a) and caecal tonsil tissue (b) of experimental chickens according to *C. jejuni* challenge status at 14 days post infection. Error bars represent IQR of the median value. Statistical significance was determined on 40 – ^Δ^Ct data using Mann Whitney-U analysis with levels of significance given as * p < 0.05, **p < 0.01, ***p < 0.001

**Figure 4.**
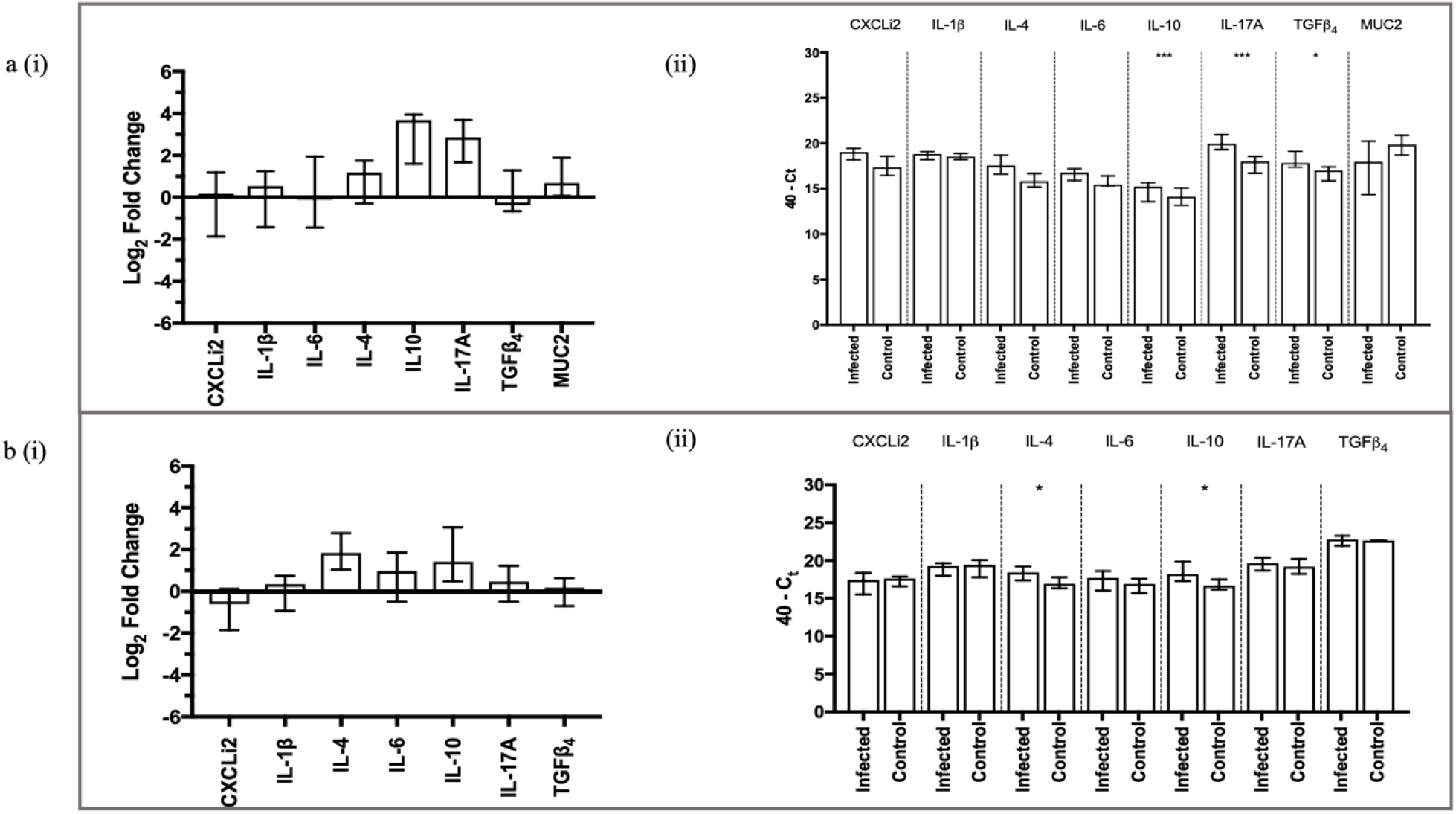
Cytokine and Mucin expression 28 days post infection. Relative expression of infected v control (i) and 40 - ^Δ^Ct (ii) of cytokine, chemokine and AMP transcripts within caecal tissue (a) and caecal tonsil tissue (b) of experimental chickens according to *C. jejuni* challenge status at 28 days post infection. Error bars represent IQR of the median value. Statistical significance was determined on 40 – ^Δ^Ct data using Mann Whitney-U analysis with levels of significance given as * p < 0.05, **p < 0.01, ***p < 0.001

**Figure 5.**
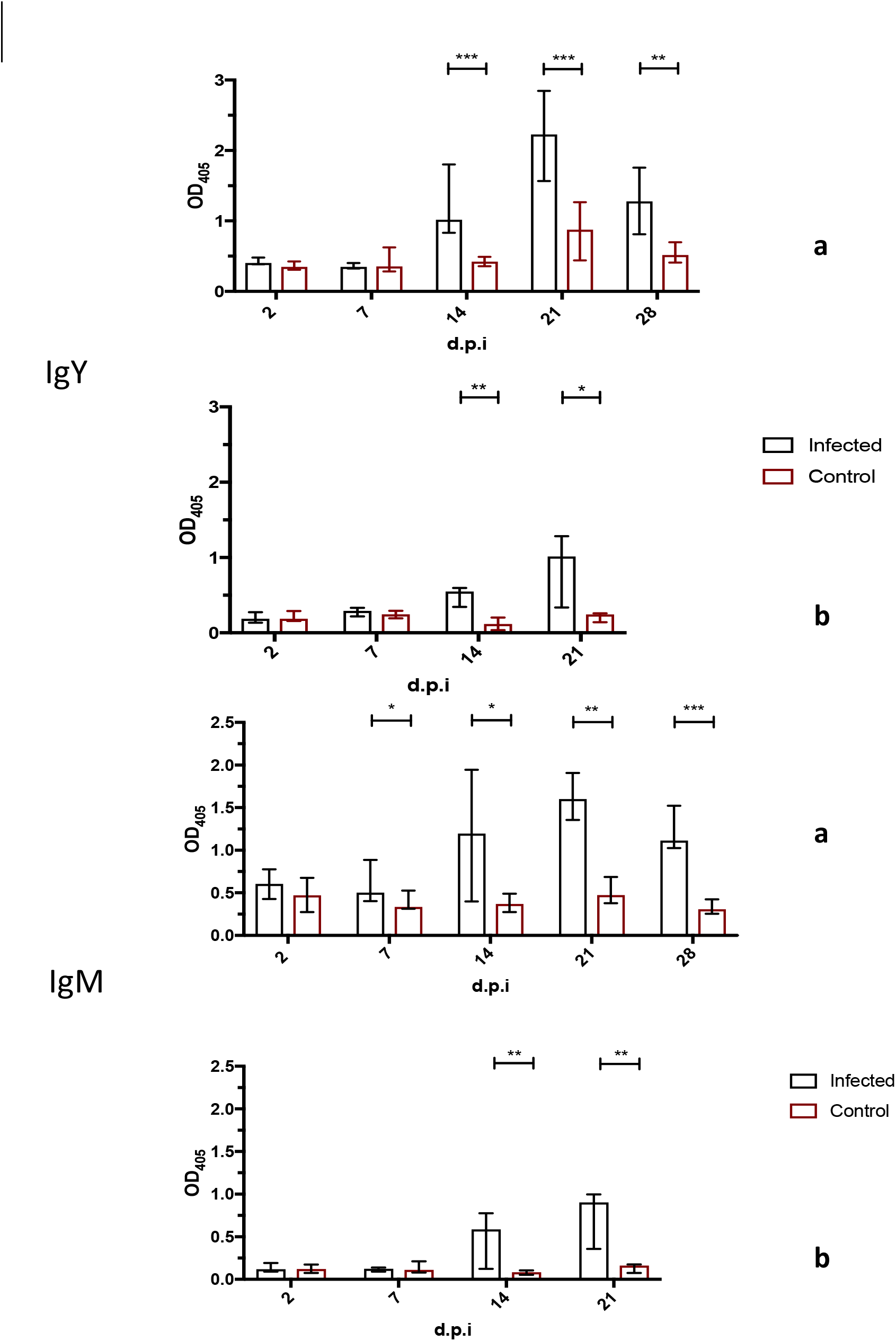
Serum IgY and IgM specific anti-*Campylobacter* antibody responses determined by ELISA following challenge with *C. jejuni* M1. **Levels of secretory IgA remain high in the gut during later infection.**

**Figure 6.**
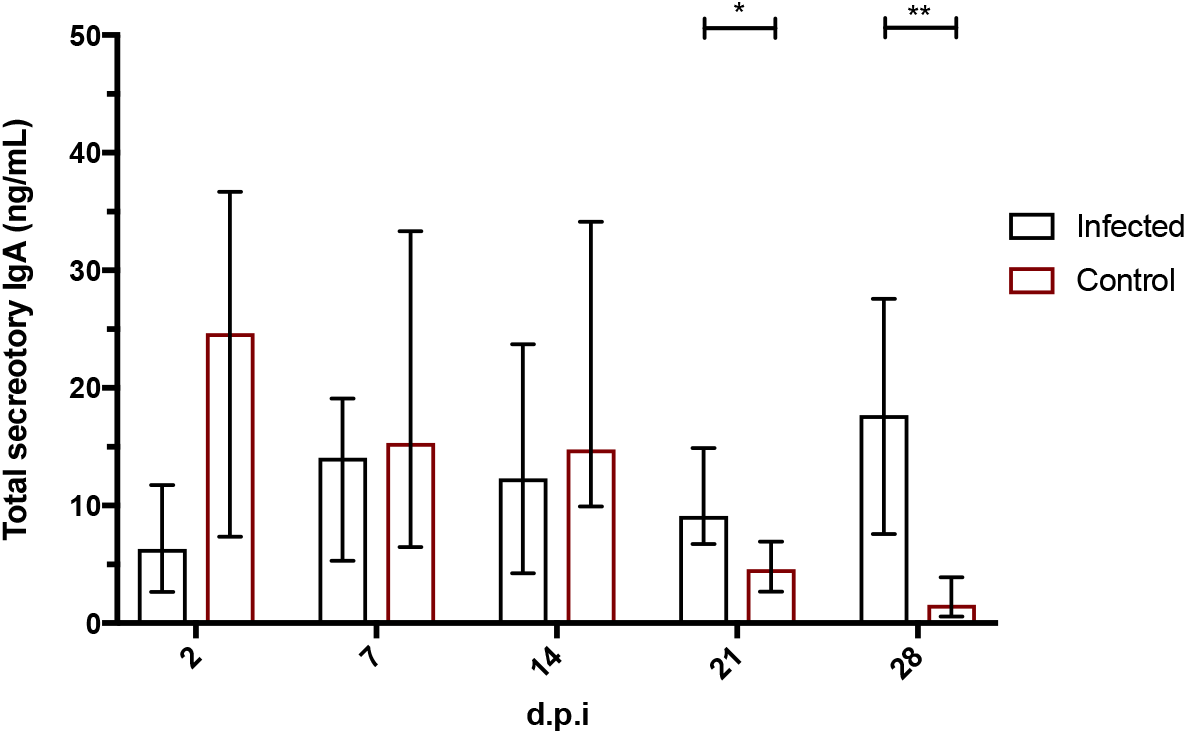
Total Secretory IgA in intestinal gut wash samples. Total secretory IgA response within ileal tissue according to *C. jejuni* M1 infection status at sampled time-points post-challenge according to experimental protocols for experiment 1 (a) and experiment 2 (b). Statistical analysis is based on median values with associated IQR. Statistical significance was determined using Mann Whitney-U analysis with levels of significance given as *p < 0.05, **p < 0.01, ***p < 0.001, ****p < 0.0001.

Taken together these data are consistent with downregulation of inflammatory responses despite *Campylobacter* persistence in the caeca, but consistent with both regulation of inflammation and the maintenance of the gut barrier. The expression of IL-4 and IL-10 in a key gut lymphoid tissue, the caecal tonsil, is consistent with maintaining a Th2 or humoral response within the gut.

### Production of specific-IgY antibody remains consistent in infection along with IL-4 expression in the caecal tonsils

Specific antibody responses against *C. jejuni* as measured in serum are consistent with a specific antibody response. By 14 days post infection significant IgM and IgY responses can be detected and remain high for the duration of the experiment.

## Discussion

Infection of the chicken intestinal tract with *C. jejuni* leads to an immune response that allows persistence in the gut but prevents invasive infection and controls inflammation. *C. jejuni* infection leads to a strong specific antibody response, though as we have previously shown, appears to have a limited role in clearance from the gut. Here we show that whilst expression of pro-inflammatory cytokines and chemokines wanes, significant levels of expression of the regulatory cytokine IL-10 and IL-17A persists within the caeca for at least 28 days following infection. In the caecal tonsils expression of IL-4 and IL-17 are maintained. We propose that the nature of the immune response to *C.jeuni* infection is one that allows persistence in the gut, but acts to limit inflammatory responses and prevent invasive infection from the gut through maintaining the intestinal barrier and secretory antibody. This ‘stalemate’ largely protects the birds from campylobacteriosis, but the persistence in the gut for the production lifetime of chickens remains a currently insoluble public health issue.

Previously we have shown that high levels of antibody persist in long term infection, but its role in clearance from the caeca is limited. We have also modelled the network of cytokine responses in infection which has proposed a key role for Th17 responses(9). The data we present here show that prolonged IL-10 and Th17 expression remain for at least 28 days post challenge. Whilst persistence of *Campylobacter* is likely to remain a driver for pro-inflammatory signals, the continued expression of IL-10 may act to control this preventing any inflammation-associated pathology. It is noteworthy that in comparison to *Salmonella* Typhimurium challenge, the magnitude of inflammatory responses is lower in magnitude but more prolonged (14-16). Induction of inflammatory responses by *C. jejuni* may help in establishing gut colonisation, by reducing hypoxia in the gut as has been proposed for *Salmonella* (17). Indeed, this may be important for a microaerophile such as *C. jejuni* in competition with anaerobic bacteria that make up the caecal microbiome of broiler chickens (18) Although Th17 responses are often associated with inflammatory conditions in mammalian species, they also play a key function in maintaining integrity of mucosal barriers through maintenance of tight junctions between epithelial cells and increased activity of key innate immune effectors such as antimicrobial peptides(11). It is unlikely that Th17 responses contribute significantly to reducing colonisation.

Indeed, increased expression of IL17A following prebiotic treatment had no effect on reducing *Campylobacter* colonisation(19), but are likely to play a key role in limiting invasive infection. It is increasingly clear from our current and previous data along with that from others that there are significant inflammatory, cellular and humoral responses to *Campylobacter* (4, 20-22), though little evidence that these are associated with caecal clearance. The idea that the chicken is ‘tolerogenic’ in its immune response is increasingly substantiated with *C. jejuni* clearly able to colonise the caeca beyond slaughter and market age, though beyond relatively mild enteritis, causing little disease in the chicken. Whilst a clear welfare issue, such inflammatory responses are rarely associated with significant morbidity let alone mortality. Thus, we propose that *C. jejuni* and the chicken are locked in a stalemate between virulent, invasive infection and the immune response. This allows *C. jejuni* to persist in the caeca, and remain a public health problem, but prevents significant levels of invasive infection and illness to the chicken host.

